# Melanoma cells appropriate pericyte:endothelial cell crosstalk during perivascular invasion in the brain

**DOI:** 10.1101/2022.07.15.500231

**Authors:** Yu Zhang, Caroline P. Riedstra, Sarah Wang, Sonia Patel, Tobias Bald, Pankaj Kumar, Ali Rohani, David F. Kashatus, Stergios J. Moschos, Andrew C. Dudley

## Abstract

Post brain colonization, cancer cells may adhere to and spread along the abluminal surface of the vasculature providing advantageous access to oxygen, nutrients, and vessel-derived paracrine factors. For example, brain-tropic melanoma cells are well-known to invade along or within blood vessels at the invasive front, but the molecular mechanisms that guide this process are not well-characterized. We have used melanoma cells of different phenotypic states (melanocytic versus mesenchymal) to characterize different modes of perivascular invasion in the brain. We find that Sox9^hi^ mesenchymal state melanoma cells undergo pericyte-like spreading along brain blood vessels via a Snail1-dependent process; in contrast, Sox10^hi^ melanocytic state melanoma form proliferative, perivascular clusters. *Snai1* deletion in mesenchymal-state melanoma cells diminishes Tgfβ-induced expression of Pdgfrβ which impairs Tgfβ/Pdgf ligand-driven motility along the brain microvasculature and dramatically reduces perivascular dispersal of melanoma cells throughout the brain post-colonization. These data suggest that, depending on their transcriptional/phenotypic state, some melanoma cells may appropriate signals that typically mediate endothelial cell:pericyte cross talk as an adaptive mechanism for maintaining vessel proximity and motility within the brain microenvironment.

## Introduction

Cancer cells invade new tissue and organ microenvironments via diverse mechanisms including single cell amoeboid-like movement and collective migration ^1-3^. Each of these mechanisms typically require cancer cells to adhere to an underlying substrate that imparts traction forces and cytoskeletal/actin remodeling to propel the cell, or clusters of cells, forward. Gradients of growth factors or other soluble substances may add directionality ^4,5^. In one mode of invasion, cancer cells “co-opt” or adhere to the abluminal surface of the vasculature (often termed angiotropism or perivascular migration, PVM) ^6,7^. For example, in the brain microenvironment, melanoma cells migrate away from the lesion core by traveling distances up to 500 µm per month while maintaining focal contacts with the perivascular surface of the brain vasculature ^8^. PVM often precedes the steps required to initiate colonization and was shown to be a contributor to dormancy, relapse, and resistance to therapy ^9,10^. However, the molecular mechanisms that guide and sustain interactions between cancer cells and the vasculature are largely uncharacterized. Given that endothelial cells are phenotypically and functionally diverse in different tissue and organ microenvironments, it is possible that cancer cells acquire equally diverse mechanisms, including changes in cell state, for interacting with and invading along the surface of the vasculature; especially in the brain/CNS where the endothelium is highly specialized.

Epithelial-to-mesenchymal transition (EMT) accompanies a cell state change that enhances cellular mobility, invasion, resistance to cell death, drug resistance, and suppression of immune surveillance ^11,12^. *Snail* genes (*Snai1*/*Snai2*) encode for potent EMT transcription factors and Snail1 expression in melanoma cells drives an immunosuppressive tumor microenvironment and partial EMT-like state ^13,14^. Snail1 is a well-characterized transcriptional repressor of cadherins, and there are important links between a Snail1-mediated mesenchymal program and invasion/drug resistance in melanoma ^15,16^. Mesenchymal-like/EMT pathways are also enriched in mesenchymal-state melanoma cells which are undifferentiated with low MITF but high AP-1 activity, and show increased migratory potential and drug resistance ^17^. The transcription factors Sox9 and Sox10 play important roles in switching melanoma cell fate: Sox9^hi^ melanoma cells are in a highly invasive mesenchymal-like state whereas Sox10^hi^ melanoma cells are in a proliferative, melanocytic state typified by expression of tyrosinase ^18^. Sox9/Sox10 expression is almost mutually exclusive and Sox10 ablation in melanocytic-state melanoma cells de-represses Sox9 and promotes acquisition of the migratory mesenchymal phenotype ^17^.

While characterizing Sox9^hi^ versus Sox10^hi^ murine melanoma cells, we found that Sox9^hi^ melanoma cells migrate along brain blood vessels via distinct pericyte-like migratory patterns whereas Sox10^hi^ melanoma cells form rounded, perivascular clusters. Vessels occupied by Sox9^hi^ melanoma cells were also longer and more dilated compared to their Sox10^hi^ counterparts suggestive of alterations in the underlying vasculature. Bulk RNAseq of mesenchymal state versus melanocytic state murine melanoma cells identified enrichment for TGFβ signaling and EMT-inducing transcription factors including *Snai1*. As proof-of-principle, we sought to address how Snail1 may control melanoma cell motility and spreading along the brain microvasculature.

## Results

### Invading melanoma cells interface with the brain microvasculature at the invasive front with distinct phenotypes

Angiotropic patterns of growth by brain-colonizing melanoma cells have been long-noted in human post-mortem biopsies ^8,19^. These angiotropic patterns of growth, either present in the primary lesion or in a metastatic site such as the brain, are typically associated with a worse overall prognosis including shorter survival. To confirm the existence of angiotropism/PVM in human brain metastasis, we examined several bioarchived specimens provided by the UVA tissue biorepository core facility. Indeed, we found numerous S100^+^ melanoma cell clusters growing along, or around CD31^+^/PECAM1^+^ brain blood vessels (Fig. 1A). While these data confirm the existence of vessel-associated melanoma cells in human brain tissues, we sought to gain insight into the molecular mechanisms that drive perivascular invasion by brain-resident melanoma cells at the earliest phases post-colonization. To do so, we used an established mouse melanoma brain metastasis model that includes: mCherry^+^ D4M (isolated from *Tyr-Cre*^*ERT2*^*;BRAF*^*V600E*^*;Pten*^*fl/fl*^ mice), mCherry^+^ Hcmel12, or tdTomato^+^ B16F10 murine melanoma cells stereotaxically injected into the right striatum of syngeneic *Cdh5-Cre*^*ERT2*^*;ZSGreen*^*l/s/l*^ mice, in which endothelial cells (ECs) are labelled with ZSGreen and herein called EC^iZSGreen^ mice.

**Fig. 1.**
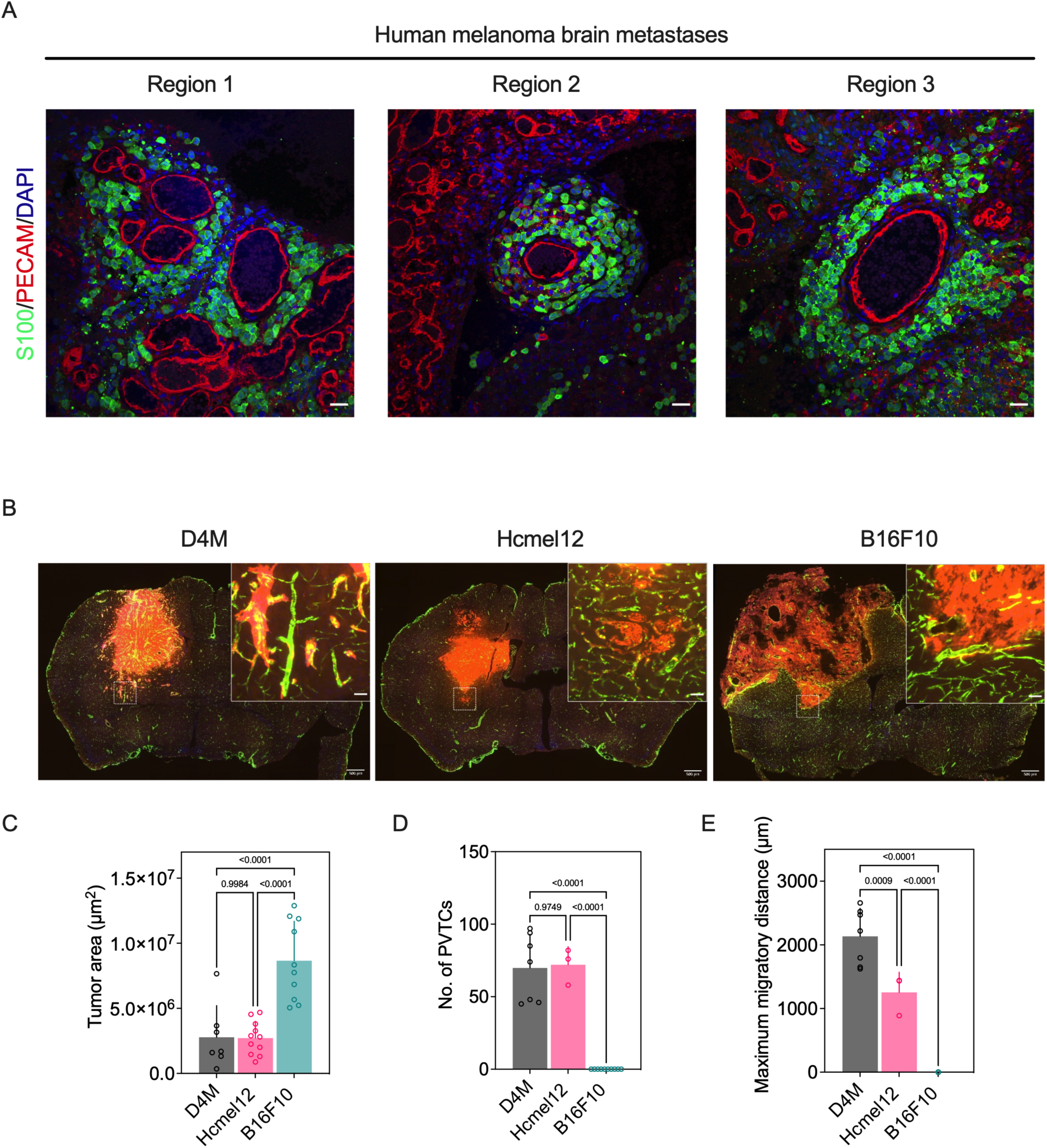
Invading melanoma cells interface with the brain microvasculature at the invasive front with distinct phenotypes. (A) Representative images of human melanoma brain metastasis samples stained with S100 and PECAM1 antibodies. (B) Representative images of brain tissue from EC^iZSGreen^ mice that were intracranially injected with mCherry^+^ D4M, mCherry^+^ Hcmel12 cells, or tdTomato^+^ B16F10 murine melanoma cells. (C) Manual quantification of bulk tumor area, number of PVTCs from the invasion front, and maximum cell cluster migratory distance. D4M; n = 4 mice, Hcmel12; n = 3 mice, and B16F10; n = 3 mice. Data are averages ± SD. All p values were determined by ordinary one-way ANOVA. Scale bars, 100 μm (A), 500 μm (B), 50 μm (insert, B).

Two weeks post-injection, tumor growth was detected by fluorescent imaging of 40 µm cryosections of the mouse brain. A large bulk tumor was observed at the striatum and cortex in all mice injected with the three melanoma cell lines, however, their invasive fronts were strikingly different. D4M cells were highly invasive with pericyte-like spreading along the abluminal surface of ZSGreen^+^ brain ECs. Although Hcmel12 cells co-opted brain vessels during invasion of the peri-tumoral environment as described previously ^20^, there was little evidence of pericyte-like spreading. By contrast, B16F10 cells formed a large, amorphous mass but did not follow similar patterns of invasion compared to the other two cells lines (Fig. 1B). Quantification of tumor area showed that B16F10 tumors were on average ∼ 2-3 fold larger compared to D4M or Hcmel12 counterparts (Fig. 1C). However, both D4M and Hcmel12 tumors showed an ∼ 50-fold increase in the numbers of perivascular tumor cells (PVTCs) and an ∼ 1,000-2,000 fold increase in maximum migratory distance (defined as the distance travelled by solitary melanoma cells or clusters of melanoma cells from the center of the bulk tumor mass) compared to B16F10 tumors (Fig. 1D and 1E). These data suggest that melanoma cells have differential abilities to engage with the brain microvasculature and perhaps disperse throughout the brain via distinct vascular-mediated mechanisms.

### D4M melanoma cells spread along the brain vasculature similar to pericytes whereas Hcmel12 form rounded, perivascular clusters

To assess the interactions between melanoma cells and the brain vasculature *in vivo*, we developed a computational analysis tool, VeCo (Vessel Co-option), which enables whole-brain mapping as well as automatic, standardized, and unbiased quantitative analysis of vessel lengths and widths in different regions of interest (ROIs). VeCo can be applied at the tumor bulk position, contralateral striatum region where no tumor cells are present, and throughout the entire remaining brain by use of whole tissue slice microscopy images. VeCo is made freely available (https://github.com/Mitogenie/VeCo). In brief, VeCo quantifies co-option by (I) extracting cancer cell or blood vessel features from fluorescence microscopy images, (II) identifying overlapping regions between cancer cells and blood vessels, (III) counting co-opted versus non co-opted cancer cells and vessels, and (IV) characterizing vessel distribution, length, and width (Fig. S1 A-F). Notably, VeCo analysis of individual vessel segments throughout entire brain sections revealed that vessels occupied by perivascular D4M cells are typically longer and wider than vessels within Hcmel12 tumors, which more closely resemble normal brain micro-vessels (Fig. 2A and 2B). D4M-occupied vessels were also ∼ 2-3 times as dilated when compared to Hcmel12-occupied vessels. No D4M cells were identified juxtaposed to vessels smaller than 7 µm; however, occasional Hcmel12 cells were found adjacent to smaller vessels. Furthermore, Hcmel12 cells were evenly distributed along vessels ranging from 30 µm to 45 µm in length whereas D4M-occupied vessels were greater than or equal to 35 µm long (a 2-3 fold increase compared to Hcmel12 cells). These data suggest that either D4M-occupied vessels are induced to become wider and longer, or that D4M melanoma cells preferentially occupy wider and longer vessels when compared to Hcmel12 counterparts. To further examine the perivascular localization of melanoma cells in relationship to pericytes, *Pdgfrb-P2A-Cre*^*ERT2*^*;ZSGreen*^*l/s/l*^ (Pericyte^iZSGreen^) mice were intracranially injected with mCherry^+^ D4M or Hcmel12 cells. D4M cells were almost exclusively spread along the abluminal surface of CD31^+^ brain endothelial cells, interdigitating with pericytes and exhibiting a pericyte-like morphology whereas Hcmel12 cells resembled “beads on a string” of rounded, perivascular cell clusters (Fig. 2C). These data suggest that melanoma cells interact with the brain microvasculature via distinct mechanisms and, by doing so, could differentially impact the function of the underlying vasculature.

**Fig. 2.**
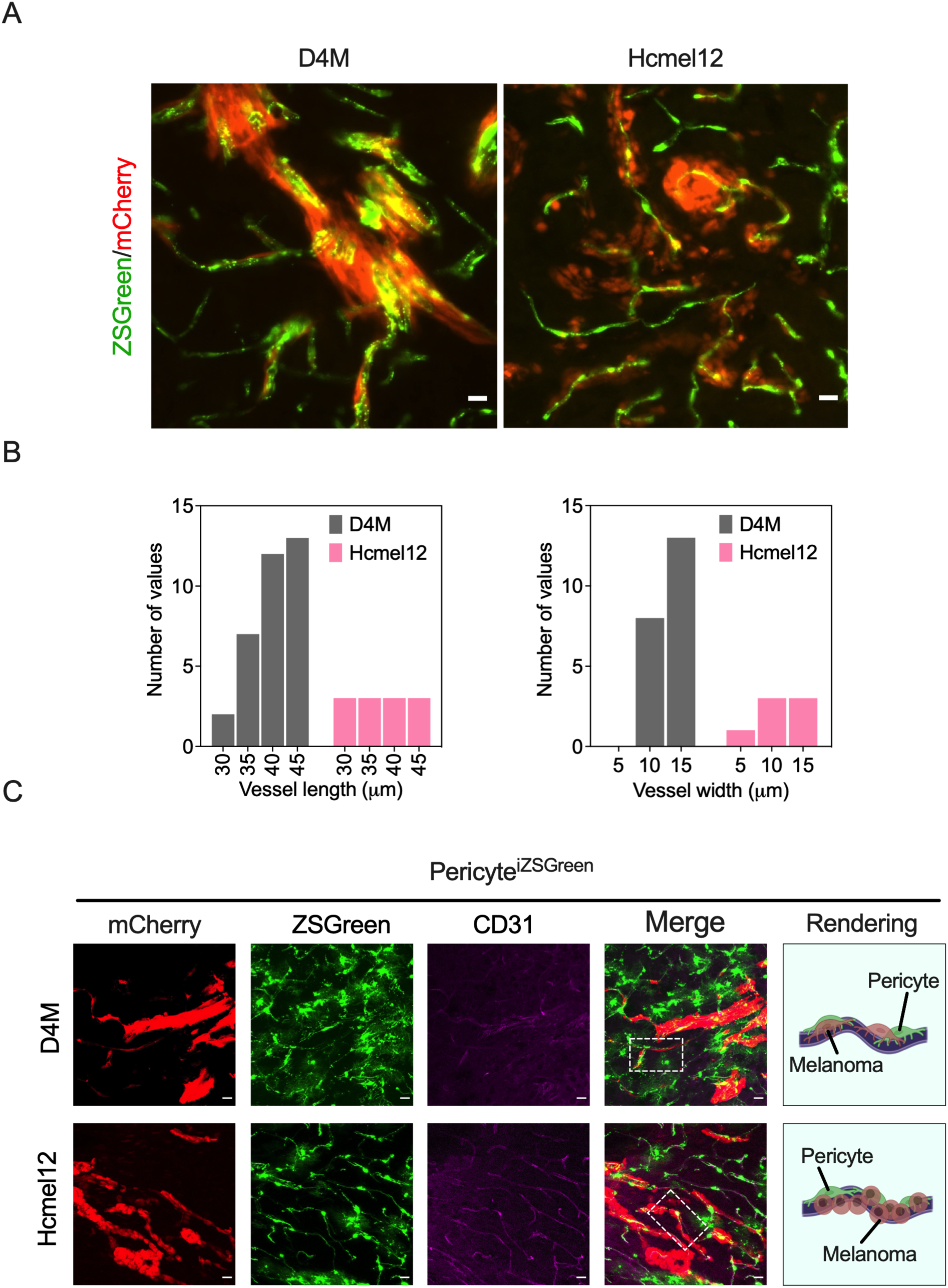
D4M melanoma cells spread along the brain vasculature similar to pericytes whereas Hcmel12 form rounded, perivascular clusters. (A) Representative images of brain tissues from EC^iZSGreen^ mice that were intracranially injected with mCherry^+^ D4M or Hcmel12 cells (representative region from the invasive front is shown). (B) VeCo analysis of co-opted vessel length and width from EC^iZSGreen^ mice injected with D4M or Hcmel12 cells. D4M; n = 4 mice, Hcmel12; n = 3 mice. (C) Representative images of brain tissue from Pericyte^iZSGreen^ (Pericyte^iZSGreen^) mice that were intracranially injected with mCherry^+^ D4M or Hcmel12 cells. DM cells are spread and interdigitated with pericytes unlike Hcmel12 which are more rounded and resemble “beads on a string” as they surround the endothelium. Scale bars, 20 μm (A and C).

### D4M cells resemble Sox9^hi^/Sox10^lo^ mesenchymal state melanoma cells

Melanoma cells exist in mutable, transitional states termed melanocytic, intermediate, and mesenchymal-like ^17,21,22^. The transcription factors Sox9 and Sox10 play binary roles in the maintenance of melanoma cell fate ^23^. Sox9^hi^ melanoma cells are in a highly invasive and drug-resistant mesenchymal-like state whereas Sox10^hi^ melanoma cells are in a proliferative, melanocytic state typified by expression of MITF ^18^ (Fig. 3A). Within this paradigm, we assessed the mesenchymal versus melanocytic signatures in D4M, B16F10, and Hcmel12 cells using qPCR and immunoblotting. At the mRNA level, we found that D4M cells express abundant known EMT/mesenchymal factors including *Snai1, Snai2, Cspg4, Pdgfrb, Fn1, Vim, Acta2*, and *Serpine1* relative to B16F10 and Hcmel12 cells (Fig. 3B). By contrast, B16F10 and Hcmel12 cells expressed the melanocytic genes *Cdh1, Tyr*, and *Mitf* which were almost absent in D4M (Fig. 3C). By immunoblotting, we confirmed that D4M were low in Sox10 expression but expressed abundant Sox9; moreover, D4M expressed abundant mesenchymal state markers at the protein level including Cspg4, Fn1, Pdgfrβ, Sma, and Snail1 which were mostly absent in B16F10 and Hcmel12 (Fig. 3D).

**Fig. 3.**
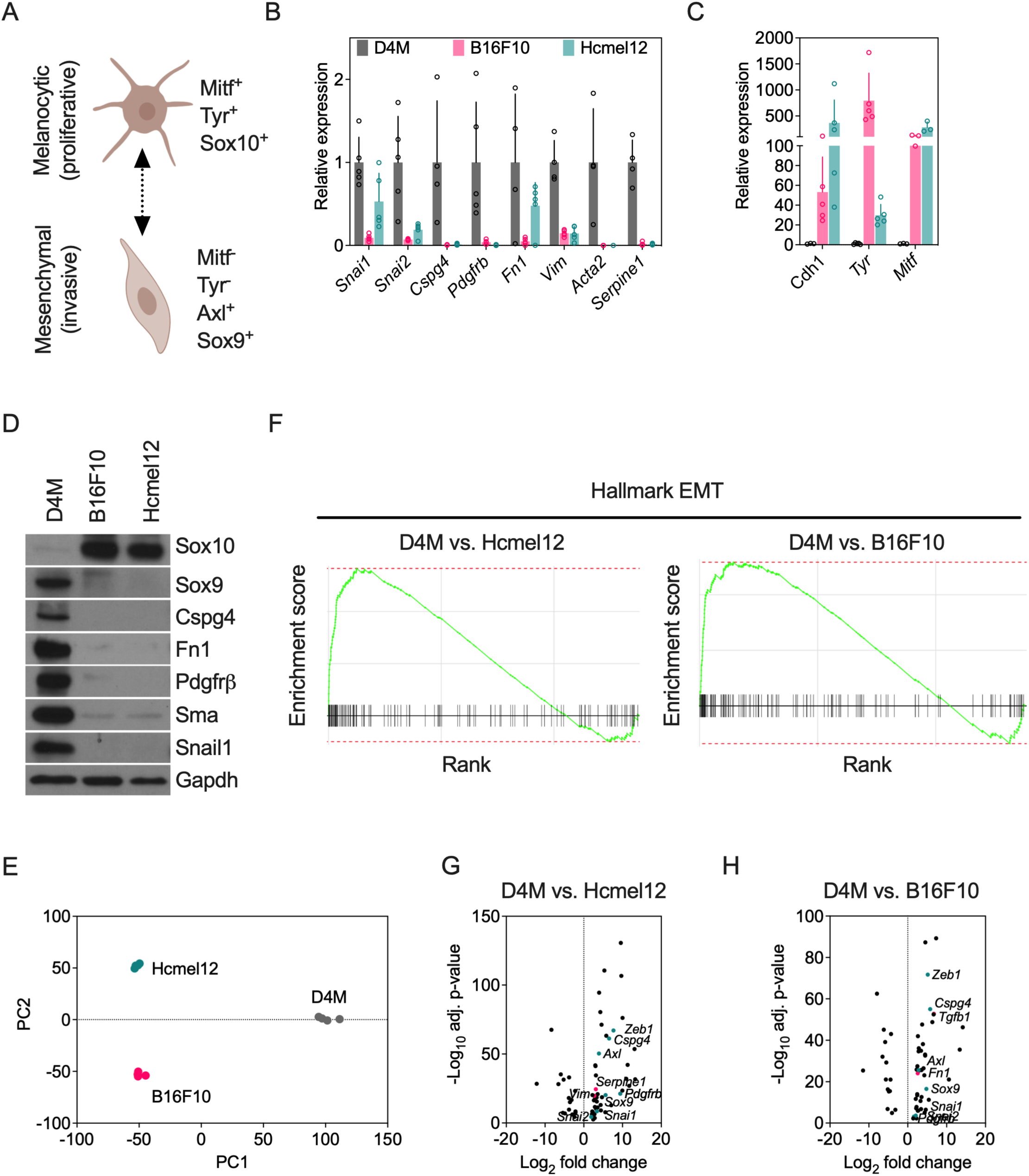
D4M cells resemble Sox9^hi^/Sox10^lo^ mesenchymal state melanoma cells. (A) Schematic of melanoma cells within different transcriptional states and their gene expression patterns. (B) qPCR examination of mesenchymal gene expression in D4M, Hcmel12, and B16F10 cells. Data are averages ± SD; n = 5 independent experiments. (C) qPCR examination of epithelial and melanocytic gene expression in D4M, Hcmel12 and B16F10 cells. Data are averages ± SD; n = at least 3 independent experiments (color legend is same as in panel B). (D) Protein expression of melanocytic and mesenchymal state genes in D4M, Hcmel12, and B16F10 cells; n = at least 3 independent experiments. (E) Principle component analyses of gene expression profile from RNAseq data from D4M, Hcmel12, and B16F10 cells; n = 4 independent samples of each cell line. (F) Gene Set Enrichment Analysis (GSEA) of EMT signature genes. (G and H) EMT signature gene expression in the indicated contrasts. Candidate EMT signature genes are depicted in pink, other genes of interest are depicted in teal. For D4M vs. Hcmel12; NES=2.45 and padj=0.009. For D4M vs. B16F10; NES=2.29 and padj=0.01.

To comprehensively characterize gene expression patterns amongst these three melanoma cell lines, we carried out bulk RNAseq. Quadruplicate samples were cleanly differentiated based on principle component analyses which separated each melanoma line by their distinct genetic signatures (Fig. 3E). Gene Set Enrichment Analysis (GSEA) confirmed that D4M cells are enriched in EMT signature genes based on normalized enrichment scores (Fig. 3F). Volcano plots of EMT signature genes, in addition to candidate mesenchymal state genes confirmed that D4M are enriched in, for example, *Zeb1, Cspg4, Axl, Serpine1, Pdgfrb, Sox9, Snai1*, and *Snai2* when compared to Hcmel12 or B16F10 (Fig. 3G and 3H).

### *Snai1* KO reduces perivascular spreading along with brain vasculature without impacting *in vitro* proliferation

As a master EMT transcription factor and transcriptional repressor of cadherins, *Snai1* (gene name = *Snai1* and protein product = Snail1) is well-characterized in breast cancer and has been shown to confer pericyte-like properties in cancer cells ^24^. Two recent studies reported an important link between a Snail1-mediated mesenchymal program and invasion/drug resistance in melanoma, but there is no known link to perivascular modes of invasion in the CNS ^15,16^. The products of *Snail* genes are potent inducers of cell movement/survival and previous work demonstrates that Snail1 expression in melanoma drives an immunosuppressive tumor microenvironment and partial EMT-like state ^13,14^. Since the functions of Snail1 in melanoma cells specifically related to perivascular mechanisms of invasion are not known, we used CRISPR/Cas9-mediated deletion of *Snai1* in D4M melanoma cells to assess its role in regulating perivascular invasion (Fig. 4A). Relative to a non-targeting sgRNA control, *Snai1* knockout (KO) had no impact on the rate of proliferation/viability *in vitro* (Fig. 4B). However, *Snai1* KO cells showed significantly reduced motility in an *in vitro* wound closure assay (Fig. 4C). Next, we intracranially injected *Snai1* KO mCherry^+^ Luciferase^+^ D4M cells (D4M^mCherry(Luc)^) into the brains of EC^iZSGreen^ mice. Tumor growth was monitored by *in vivo* bioluminescence imaging (BLI) over the course of two weeks. The results showed that *Snai1* KO strikingly reduced D4M growth in the brain resulting in a five-fold reduction in photon flux at 14 days after tumor cell implantation (Fig. 4D). Fresh cryosections prepared at the end of the experiment confirmed a dramatic reduction in bulk tumor area with qualitatively less evidence of perivascular invasion and fewer solitary “satellite” melanoma cells appearing at the invasive front (Fig. 4E). Using ImageJ analysis from these cryosections, we calculated that bulk tumor area was reduced by ∼ 90% in *Snai1 KO* cells when compared to the control D4M cells (Fig. 4F). By focusing on the bulk tumor margins (invasive front), we found an ∼ 80% reduction in PVTCs in *Snai1* KO D4M cells relative to control; furthermore, those *Snai1* KO cells that did occupy the surrounding vasculature were smaller and showed less evidence of perivascular spreading compared to their wildtype counterparts (Fig. 4G and 4H). Taken together, these data suggest that *Snai1* deletion in melanoma cells profoundly impairs tumor expansion and perivascular invasion in the brain *in vivo* despite no evidence of an inhibitory effect on *in vitro* proliferation and a modest effect on *in vitro* migration/invasion.

**Fig. 4.**
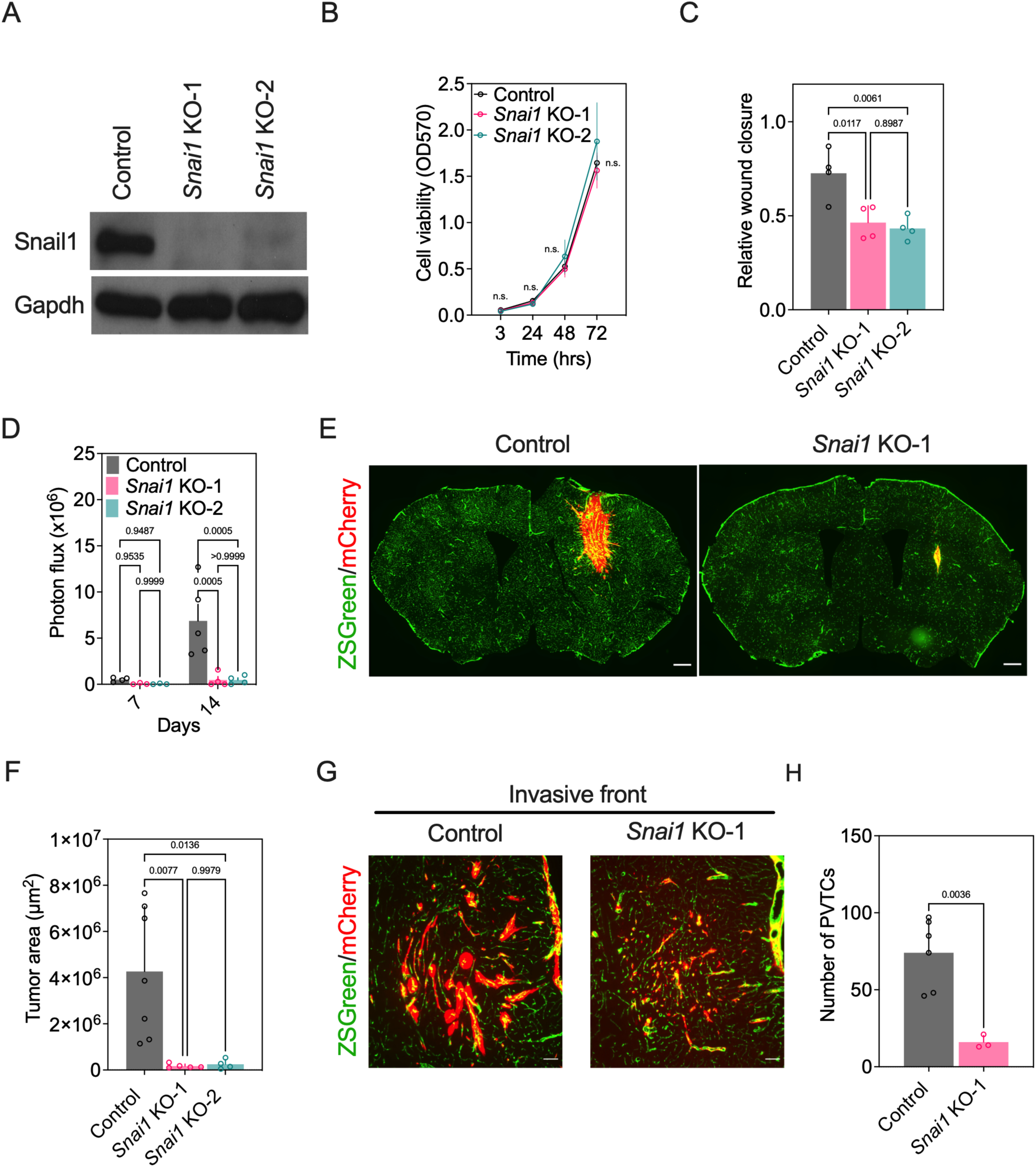
*Snai1* KO reduces perivascular spreading along with brain vasculature without impacting *in vitro* proliferation. (A) Snail1 protein expression in D4M non-targeting control or *Snai1* KO cells; n = 3 independent experiments. (B) Cell viability examined by crystal violet assay; n = 3 independent experiments. (C) Cell motility examined by *in vitro* wound closure assay; n = 3 independent experiments. (D) Photon flux in EC^iZSGreen^ mice injected with mCherry^+^ Luciferase^+^ D4M control or *Snai1* KO cells; control, n = 5 mice and *Snai1* KO, n = 4 mice. (E) Representative images of brain sections from EC^iZSGreen^ mice injected with mCherry^+^ D4M control or *Snai1* KO cells. (F) Quantification of bulk tumor area from EC^iZSGreen^ mice injected with mCherry^+^ D4M control or *Snai1* KO cells; control, n = 7 mice, *Snai1* KO-1, n = 5 mice, *Snai1* KO-2, n = 4 mice. (G) Representative images of the invasive front in EC^iZSGreen^ mice injected with mCherry^+^ D4M control or *Snai1* KO cells. (H) Number of perivascular melanoma cell clusters at the invasive front in EC^iZSGreen^ mice injected with mCherry^+^ D4M control or *Snai1* KO cells; control, n = 6 mice, *Snai1* KO-1, n = 3 mice. Scale bar, 500 μm (E), 50 μm (G). P values were determined by two-way ANOVA (B and D), Student’s t-test (C and H) and one-way ANOVA (F).

### *Snai1* deletion impairs the motility of D4M cells along brain microvessels by disruption of a TGFβ/Pdgfb/Pdgfrβ signaling axis

Next, we used an *ex vivo* brain slice culture model to quantify perivascular motility by D4M cells -/+ *Snai1* deletion (Fig. 5A) ^25,26^. In this assay, mCherry^+^ melanoma cells were seeded on a 250 µm thick, freshly prepared brain slice from EC^iZSGreen^ mice. Following a 2∼3 day incubation, live imaging was carried out to trace the perivascular migration of individual melanoma cells. *Snai1* KO reduced the motility of melanoma cells along the brain vasculature by ∼ 50% relative to control cells, suggesting that Snail1 is important for regulating perivascular spreading and motility (Fig. 5B and 5C). To examine whether Snail1 is sufficient to promote perivascular spreading and tumor expansion, melanocytic state B16F10 cells with stable *Snai1* overexpression (OE) were used for brain slice culture assays and brain tumor injections (Fig. S2A). Consistent with *Snai1*’s ability to promote a partial EMT-like state, Snail1 OE was sufficient to up-regulate Pdgfrβ, Fn1, and Sma expression in B16F10 melanoma cells (Fig. S2A). Compared with B16F10 control cells, *Snai1* OE cells exhibited increased migratory velocity (∼2-fold) along the vasculature in the *ex vivo* model (Fig. 5D). Two weeks after intracranial injection, *Snai1* OE cells also showed increased tumor area inside the brain cortex, and greater numbers of cells spreading in diffuse patterns along brain blood vessels at the invasive front (Fig. S2B and S2C). This is contrasted with B16F10 control melanoma cells which instead were more rounded in appearance and grew in proliferative clusters around, rather than along pre-exiting vessels.

**Fig. 5.**
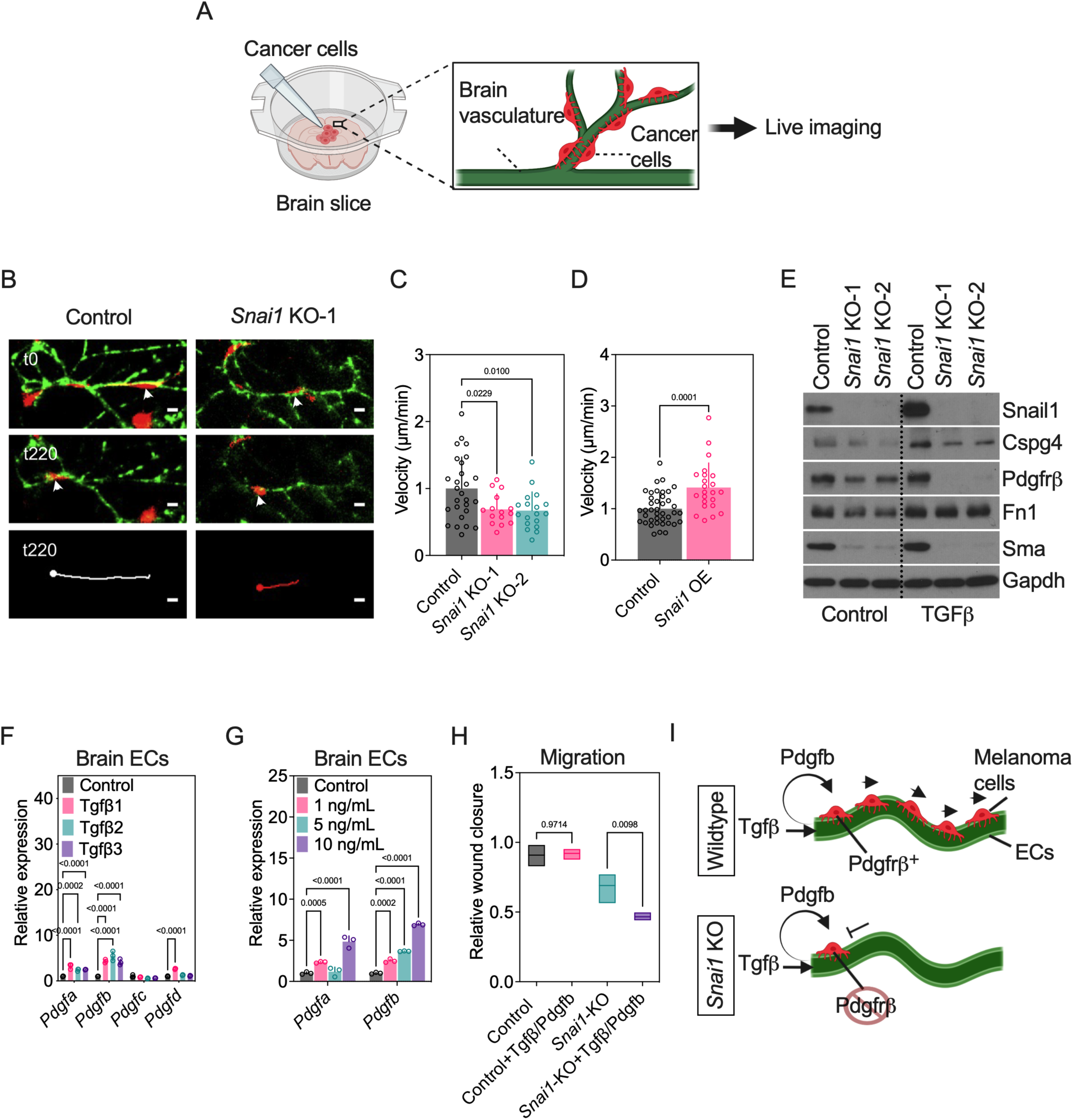
*Snai1* deletion impairs the motility of D4M cells along brain microvessels by disruption of a TGFβ/Pdgfb/Pdgfrβ signaling axis. (A) Schematic of brain slice and tumor cell culture model. (B) Representative images of mCherry^+^ D4M control or *Snai1* KO cells spreading and migrating along ZSGreen^+^ brain vessels in the *ex vivo* brain slice culture model. (C) Quantification of migratory velocity of D4M control or *Snai1* KO cells; n = 3 independent experiments. (D) Quantification of migratory velocity of B16F10 control or *Snai1*-OE cells; n = 3 independent experiments. (E) Protein expression in D4M control or *Snai1* KO cells with/without TGFβ1 treatment; n = 3 independent experiments. (F) *Pdgf* mRNA levesl in BrECs cells treated with/without Tgfβ1, Tgfβ2, or Tgfβ3; n = 3 independent experiments. (G) *Pdgf* mRNA levels in BrECs treated with different doses of Tgfβ2; n = 3 independent experiments. (H) Wound closure assay of D4M control or *Snai1* KO cells treated with/without the combination of Tgfβ1 (10 ng/mL) and Pdgfb (10 ng/mL); n = 3 independent experiments. (I) A model for Snail1-mediated perivascular spreading of melanoma cells and possible crosstalk with BrECs through a Tgfβ/Pdgfb/Pdgfr signaling axis during perivascular dispersal in the brain. Scale bar, 20 μm (B). P values were determined by one-way ANOVA (C and H), Student’s t-test (D), or two-way ANOVA (F and G).

To investigate the mechanism whereby Snail1 mediates the perivascular invasion of melanoma cells, we first examined expression of various mesenchymal marker proteins (Cspg4, Pdgfrβ, Fn1, and Sma) in control versus *Snai1* KO cells, and found *Snai1* deletion reduced the expression of these candidate genes to varying levels (Fig. 5E). Since Tgfβ is a well-known inducer of EMT and mesenchymal state genetic programs in general, we asked whether *Snai1* KO would impact the expression of these candidate factors upon exposure to Tgfβ1. The results showed that most of these genes were unchanged or partially reduced by Tgfβ1 in the absence of *Snai1*; surprisingly, however, we found in the presence of Tgfβ1, *Snai1* KO cells showed a further reduction in Pdgfrβ expression (Fig. 5E).

It is well-established that endothelial cell-secreted Pdgfb is critical for the recruitment of Pdgfrβ^+^ pericytes towards blood vessels ^27,28^, that Tgfβ induces differentiation of mesenchymal cells into pericytes ^29^, and that Pdgf and Pdgfrβ are upregulated by Tgfβ stimulation ^30^; thus, we hypothesized that mesenchymal-state melanoma may appropriate a Tgfβ/Pdgf/Pdgfrβ signaling axis to facilitate crosstalk with brain endothelial cells (BrECs) during perivascular spreading and invasion. Indeed, we found after treatment with Tgfβ ligands, cultured murine BrECs showed increased *Pdgfa* and *Pdgfb* levels in a ligand-dependent and dose-dependent manner, with *Pdgfb* showing the most robust up-regulation (Fig. 5F and 5G). Furthermore, whereas control cells showed enhanced migration at six hours when treated with combinations of Tgfβ1 and Pdgfb ligands, there was no further increase in migration at the 12 hour time point once the cells had reached confluency; in contrast, *Snai1* KO cells showed significant impairment in cell motility in wound closure assays at 12 hours when compared with *Snai1* KO cells without treatment (Fig. 5H and data not shown). Taken together, these data demonstrate a Snail1-mediated perivascular spreading and expansion of mesenchymal state melanoma cells in the brain and suggests that crosstalk between melanoma cells and BrECs through a Tgfβ/Pdgfb/Pdgfrβ signaling axis facilitates perivascular invasion and/or retention of melanoma cells along the abluminal surface of the endothelium (Fig. 5I).

## Discussion

Identifying the fundamental molecular mechanisms that enable perivascular spreading and growth of cancer cells in the brain microenvironment will guide the development of new therapeutic approaches to prevent brain colonization, dormancy, and spreading throughout the brain. Here we show that, depending on cell state (melanocytic versus mesenchymal), melanoma cells differentially occupy perivascular niches and interface with the vasculature with distinct phenotypes. Specifically, we demonstrate Snail1-mediated perivascular spreading and expansion of mesenchymal-state melanoma cells and suggest an important role for a Tgfβ/Pdgfb/Pdgfrβ axis in guiding and sustaining pericyte-like spreading and growth of melanoma cells in the brain post-colonization. Thus, the transcriptional state of the melanoma cells themselves is an important determinant of PVM and a complimentary mechanism of vessel-mediated dispersal alongside previously hypothesized mechanisms including chemotactic soluble factors (bradykinin, CXCR4-binding cytokine and Wnt 7a/b), adhesion molecules (β1 integrin, α6 integrin and L1CAM), and actin-related protein complex Arp2/3 ^9,25,31-35^.

Post brain colonization, cancer cells may acquire new phenotypic or metabolic dependencies (e.g. lipid metabolism and oxidative phosphorylation) or resurrect developmental transcriptional programs that enable their survival and invasion ^36^. For example, melanoma cells have been shown to re-instate neural crest-specific transcription factors that may provide an adaptive, survival advantage in the brain microenvironment ^16,37-39^. Notably, while one recent study has found that AXL and EMT-like melanoma cells are enriched in melanoma brain metastases, another recent study found that brain metastases were instead enriched in neuronal-lineage genes and melanocytic state markers such as MITF and SOX10 ^40,41^. Most likely, melanoma phenotypic plasticity produces a spectrum of different lineage states that interact cooperatively to maintain tumor survival, persistence, and invasion as they compete for various niches in the brain microenvironment (i.e., perivascular positioning).

Angiotropic melanoma cells can also be observed at primary sites (i.e. the dermis) which, along with primary tumor ulceration, is associated with worse overall survival ^20^. Historically, associations between clinical outcomes and angiotropism in the brain have been challenging to establish, often due to small patient sample size, tissue necrosis, or the use of small amounts of biopsied material which makes it challenging to quantify perivascular invasion specifically along the invasive front ^42^. To begin to address these challenges, we used murine models of “established” brain metastasis so that we could study the behavior of incipient melanoma cells emigrating outward from a single lesion focus in mice with genetically labelled vasculature. Since there are few models of spontaneous melanoma brain metastasis in mice, future studies may expand this work to include more physiologically relevant models of brain seeding which includes intracardiac injections of melanoma cells within different transcriptional states.

In the present study, we refined our focus to include consideration of melanoma heterogeneity/plasticity that has been linked to expression of the transcription factors Sox9 and Sox10. Sox9^hi^ melanoma cells are programed to express mesenchymal-lineage genes which enables their invasion while suppressing proliferation. In contrast, Sox10^hi^ melanoma cells are in a highly proliferative state, they lack expression of typical mesenchymal-lineage genes, yet they retain melanocytic markers such as Tyr and Mitf (and pigmentation). In brief, we find that D4M melanoma cells strongly resemble human Sox9^+^ mesenchymal-state melanoma cells previously defined by other laboratories ^15,17^. In contrast, B16F10 and Hcmel12 melanoma cells resemble Sox10^+^ melanocytic state melanoma cells defined by these same groups. By comparing/contrasting the behavior of these different melanoma cells in the brain post-colonization, we identified three common characteristics; namely, (1) melanocytic B16F10 cells form well-defined tumor margins composed of highly proliferative cells without evidence of perivascular invasion but with strong evidence of angiogenesis based on intratumoral vessel morphology (large dilated vessels, abundant tortuosity, and high microvascular densities), (ii) Hcmel12 cells form smaller tumors with poorly defined margins but show features of “classic” vessel co-option typified by rounded perivascular clusters without evidence of disruption of the occupied vasculature or stimulation of angiogenesis and (iii) D4M cells are almost exclusively positioned along the abluminal surface similar to pericytes and they spread and invade along the vasculature by maintaining contact with the endothelium (or the endothelial basement membranes) based on live image analysis of fresh brain slices.

Using a combination of candidate approaches and bulk RNAseq, we confirmed the mesenchymal-like state of D4M versus counterpart melanocytic state melanoma cells and identified potential transcriptional pathways that were chosen for downstream gain/loss of function studies. One candidate includes the well-known EMT transcription factor Snail1. In breast cancer and head/neck cancers, Snail1 defines a partial EMT phenotype and it plays a functional role in collective invasion ^43,44^. Using complimentary knockout and overexpression of *Snai1*, we report that Snail1 is necessary and sufficient for melanoma cells to carry out perivascular spreading and growth. Deletion of *Snai1* from mesenchymal-like D4M melanoma cells resulted in reduced perivascular spreading and strikingly reduced tumor growth in the brain. By contrast, *Snai1* overexpression in melanocytic B16F10 melanoma cells conferred partial pericyte-like spreading along brain blood vessels, resulted in a dramatic increase in morbidity, and increased intracranial tumor cell invasion and growth.

While there remains much to be determined mechanistically as it relates to the specific role for Snail1 in programming perivascular invasion by melanoma cells, our data did uncover an unexpected loss of Pdgfrβ expression in *Snai1* KO cells that were challenged with the well-known Snail1/EMT inducer Tgfβ. These data led us to hypothesize that Tgfβ-induced expression of Pdgf ligands in the endothelium, or perhaps by melanoma cells directly, might be an important driver of perivascular invasion by melanoma cells and that disruption of this pathway instigated by loss of Pdgfrβ in *Snai1* KO cells could explain our observed phenotype. A more comprehensive investigation of endothelial cell:tumor cell crosstalk via Tgfβ/Pdgfb/Pdgfrβ signaling may require, however, more complex studies using transgenic mice with endothelial cell-specific deletion of Pdgf ligands and/or corresponding deletions of Tgfβ receptors or Pdgfrβ in melanoma cells.

In conclusion, we find that *Snai1* knockout melanoma cells lose mesenchymal features, show reduced cell motility/perivascular spreading, and reduced growth in the brain. In contrast, overexpression of *Snai1* promotes mesenchymal-like features and perivascular invasion of melanoma cells. Snail1 regulates the expression of Pdgfrβ, which plays an essential role in promoting proliferation, migration, and recruitment of pericytes and smooth muscle cells towards endothelial cells. In response to Tgfβ, melanoma cell may co-opt Tgfβ/Pdgfb/Pdgfrβ signaling that mediates endothelial cell:pericyte cross talk as a mechanism for pericyte-like spreading and dispersal along the vasculature. Disrupting the specific, cohesive interactions between brain-resident melanoma cells and the vasculature may provide a therapeutic approach to reduce brain colonization and invasion.

## Methods

### Animal studies

All experiments were performed in accordance with the University of Virginia guidelines for animal handling and care. *Cdh5-Cre*^*ERT2*^*;ZSGreen*^*l/s/l*^ and *Pdgfrb-P2A-Cre*^*ERT2*^*;ZSGreen*^*l/s/l*^ mice were all on C57BL/6 background. *Cdh5-Cre*^*ERT2*^ mice were generated by Ralf Adams (Max Planck Institute for Molecular Biomedicine). *Ai6 ZsGreen* (#007906) and *Pdgfrb-P2A-Cre*^*ERT2*^ (#030201) were purchased from The Jackson Laboratory. All mice used were around 8-weeks old and fed a standard diet and co-housed with mice of the same cohort. Tamoxifen was used to induce the expression of Cre-controlled genes (e.g., the ZSGreen reporter). Mice at 6 to 7 weeks of age were treated 3 times over the course of 7 days with i.p. injections of 75 mg/kg tamoxifen. For intracranial injection, 5,000 melanoma cells in 3 µl HBSS were delivered via stereotaxic injections into the right striatum of EC^iZSGreen^ or Pericyte^iZSGreen^ mice. Tumor growth was monitored by BLI at 1 week or 2 weeks post injection, using intraperitoneal injection of 5 mM CycLuc1 (AOB1117) dissolved in 100 µl PBS and the IVIS Spectrum Xenogen instrument. Mice were anesthetized with Ketamine/Xylazine cocktail (80-100 mg/kg and 5-10 mg/kg, respectively), perfused with PBS and fixed in 4% PFA.

### Tissue processing and immunohistochemistry

Isolated mouse brain tissues were post-fixed in 4% PFA overnight at 4°C, and then dehydrated in 30% sucrose solution for 48 hours at 4°C prior to being embedded in O.C.T. and cut into 40 μm brain slice sections using a Leica microtome. Brain slices were washed in PBST (PBS + 0.2% Triton X-100) and incubated in blocking solution (PBST + 5% normal goat serum + 5% BSA) for 1 hr at room temperature. Brain slices were then incubated with rat-anti-mouse CD31 primary antibody (1:100, BD550274) in the blocking solution at 4°C overnight. Slices were washed with PBST, and then incubated with Alexa-647 goat-anti-rat secondary antibody (1:500, Invitrogen) in the blocking solution for 1 hr at room temperature. After washing, slices were mounted with Vectashield. Paraffin sections from human melanoma brain metastasis specimens were stained according to standard protocol. In brief, slices were blocked in TBS with 5% goat serum and 5% BSA, followed by incubation of primary antibody mouse-anti-human PECAM1 (1:100, CM347A, Biocare), and rabbit-anti-human S100 (1:100, CP021A, Biocare) overnight at 4°C. After secondary antibody Alexa-488 goat-anti-rabbit (1:200, Invitrogen) and MACH2 mouse AP-polymer incubation, slices were washed and developed with the Warp Red Chromogen kit, stained with DAPI, and mounted with Vectashield. Images were taken using a ZEISS LSM 880/900 with Airyscan 2 or Nikon Eclipse Ti-E inverted microscope.

### Organotypic brain slice culture

Brains from 8-week-old EC^iZSGreen^ mice were dissected in HBSS, and cut into 250 μm slices using a tissue slicer (Stoelting). Brain slices were then cultured on Millicell cell culture inserts (PICM0RG50, Millipore) in culture media, and incubated at 37°C and 5% CO_2_ for 1 hr. Twenty-five hundred mCherry^+^ D4M control*/Snai1* knockout or B16F10 control/*Snai1*-OE melanoma cells were suspended in 1 µl culture media and seeded on top of each brain slice and incubated for 48-72 hours. Confocal Z-stack time-lapse images were taken using a ZEISS LSM 880/900 with Airyscan 2 microscope over a 3-4 hour period with a 20-minute interval. Distance and velocity of individual melanoma cells with perivascular migration was quantified with the ImageJ cell tracker plugin.

### Cell culture

D4M3A cells were cultured in DMEM/F-12 medium supplemented with 5% fetal bovine serum (FBS), Antimycotic/Antibiotic, and Plasmocin. B16F10, HEK293T, and brain vascular endothelial cells (b.End3) were cultured in 4.5g/l D-glucose DMEM supplemented with 10% FBS, Antimycotic/Antibiotic, and Plasmocin. Hcmel12 cells were cultured in RPMI 1640 medium (with 2mM L-glutamine) supplemented with 10%FBS, 10mM non-essential amino acids, 1mM HEPES, Antimycotic/Antibiotic/Plasmocin, and freshly-added 2-mercaptoethanol. For Tgfβ treatment of b.End3 cells, cells were incubated with Tgfβ1, Tgfβ2, or Tgfβ3 (10 ng/mL) or Tgfβ2 (1 ng/mL, 5 ng/mL, and 10 ng/mL) in 4.5g/l D-glucose DMEM supplemented with 1% FBS for 48 hours.

### Wound closure assay

D4M control or *Snai1* knockout cells were plated at 2×10^5^ cells per well in a 0.5% gelatin coated 6-well tissue culture treated plate. Following a 24 hours incubation, the monolayer was gently scratched with a 200 μl pipette tip across the center of the well. The cells were washed once with PBS and then cultured in low serum medium with 0.5% FBS. Images were taken at 0, 6, and 12 hrs with a Nikon Eclipse Ti-E inverted microscope with NIS-Elements software package. Wound area was analyzed with ImageJ software.

For the ligand-treated wound closure assays, cells were plated on 0.5% gelatin coated 6-well tissue culture treated plates at 5×10^4^ cells per well. After an overnight incubation period to allow the cells to adhere, media was switched from standard 5% FBS DMEM/F-12 to low serum medium with 0.5% FBS. The cells were then treated with 10ng/mL of Pdgfb and 10ng/mL Tgfβ1 in combination. Cells were incubated for 48 hours prior to being scratched with a 200µL pipette tip. After scratching, the wells were rinsed with PBS and then placed into fresh low serum media containing the previously mentioned ligand concentrations. Wells were imaged at 0, 6, and 12 hrs with Nikon Eclipse Ti-E inverted microscope and NIS-Elements software package, and analyzed with ImageJ software.

### Crystal violet assay

Crystal violet assays were conducted using standard methods. In brief, D4M control or *Snai1* knockout cells were plated at 2,500 cells per well in a 96-well plate. The cells were stained with 0.5% crystal violet staining solution and incubated for 20 minutes at room temperature. After washing with tap water and drying, methanol was added to the well to dissolve the crystal violet. The optical density at 570 nm was measured with a plate reader. Each assay was conducted in triplicate, and the OD570 of three wells without cells determined the background of staining method.

### *Snai1* knockout and overexpression cells

In brief, mouse *Snai1* genomic sequence was submitted to an online sgRNA design tool (https://portals.broadinstitute.org/gppx/crispick/public) to select three target sites. Oligonucleotides for each target site were designed, annealed and then cloned into lentiCRISPRV2 vector. HEK293T cells were co-transfected with lentiCRISPRV2-*Snai1* gRNA, vsvg and delta 8.9 vectors using X-tremeGene HP DNA transfection reagent (06366244001, Sigma). Two days after transfection, HEK293T media containing lentivirus was collected, filtered through 0.2 μm filter, and added directly to mCherry^+^ D4M3A cells with polybrene (8 μg/mL). Two days after infection, Puromycin (2 μg/mL) was added to select infected cells. Then D4M3A cells were passaged, and seeded as single colonies in 96-well plates. After expansion, the expression of Snail1 in each colony was examined using western blotting. Two clones showed complete reduction of Snail1 protein expression. The efficient oligonucleotides used to construct *Snai1* sgRNA were: *Snai1* gRNA-forward (5’-caccgATGACTGGATATGAACTCCT-3’), *Snai1* gRNA-reverse (5’-aaacAGGAGTTCATATCCAGTCATc-3’). Oligonucleotides used to construct non-targeting sgRNA were: NT gRNA-forward (5’-caccgAAAACGGCTCGATCGGTGAT-3’), NT gRNA-reverse (5’-aaacATCACCGATCGAGCCGTTTTc).

To generate stable Snai1-overexpressing cells, the mouse *Snai1* gene was amplified from pTK*-Snai1* plasmid (#36976, Addgene) using primers (forward: 5’-gcgtctagagccaccATGCCGCGCTCCTTCCTG-3’; reverse: 5’-gcgggatccaggaccggggttttcttccacgtctcctgcttgctttaacagagagaagttcgtggcctcgagaccggtGCGAGGGCCTCCGGAG CA-3’), and cloned into pLV-mCherry vector (#36084, Addgene) to obtain the pLV-*Snai1*-p2A-mCherry plasmid. HEK293T cells were transfected with pLV-mCherry/pLV-*Snai1*-p2A-mCherry, psPAX2, and pMD2.G plasmids using lipofectamine 3000 reagent (L3000001, thermofisher). B16F10 cells were infected to obtain control (pLV-mCherry) or Snai1 overexpression (pLV-*Snai1*-p2A-mCherry) cells.

### Immunoblotting

Protein extracts and western blotting was carried out using standard methods. Antibodies used were NG2 (AB5320), PDGFRβ (CS4564), FN1 (AB2413), SMA (A5228), SNAI1 (CS3879), SOX9 (SC166505), SOX10 (SC365692), CDH1 (SC3195) and GAPDH (CS5174).

### Real-time quantitative PCR

RNA extraction and real-time PCR were carried using standard methods. The primers used for real-time PCR are listed in Table 1.

### RNAseq

RNA samples were harvested from cells and sent to Novogene for bulk RNA sequencing. Sequencing analysis was performed by the UVA Bioinformatics Core. On average, we received 30 million paired ends for each of the replicates. RNAseq libraries were checked for their quality using the fastqc program (http://www.bioinformatics.babraham.ac.uk/projects/fastqc/). The results from fastqc were aggregated using multiqc software. In-house developed programs were used for adaptor identification, and any contamination of adaptor sequence was removed with cutadapt (https://cutadapt.readthedocs.io/en/stable/). Reads were mapped with the “splice aware” aligner ‘STAR’ to the transcriptome and genome of mm10 genome build. The HTseq software was used to count aligned reads that map onto each gene. The count table was imported into R to perform differential gene expression analysis using the DESeq2 package. Low expressed genes (genes expressed only in a few replicates and had low counts) were excluded from the analysis before identifying differentially expressed genes. Data normalization, dispersion estimates, and model fitting (negative binomial) were carried out with the DESeq function. The log-transformed, normalized gene expression of 500 most variable genes was used to perform an unsupervised principal component analysis. The differentially expressed genes were ranked based on the log2fold change and FDR corrected p-values. The ranked file was used to perform pathway analysis using GSEA software. The enriched pathways were selected based on enrichment scores as well as normalized enrichment scores.

### Computational methods

VeCo was developed in Python and it offers a semi graphical user interface through Jupyter notebooks for ease of use. Our developed tool is optimized to analyze 2D fluorescence coronal scans from mice. The analysis process starts by separately importing the vessel and tumor channels from the brain slices. Once imported, we apply a top-hat filter to the vessel channel to reduce the effects of possible non-uniform illumination. This step is followed by signal thresholding to create a binary mask of the vessel network. After creating the mask, we consecutively apply morphological closings and openings to remove noise and fill holes created across some vessels. Finally, we use a median filter to remove unwanted noise introduced through application of the previous filters. These pre-processing steps assure that the images have optimal quality for quantification. Before quantifying vessels, we mark two regions on the brain to create three different zones: tumor (bulk tumor), control (contralateral striatum with no tumor cells), and remaining vessels. The user crops each of these regions, and the vessel quantification is performed independently within each region. To perform vessel quantification, we first extract the skeleton of the vessel network across the brain through a thinning process. In this process, we continuously erode the vessel network from outside to the point where further erosion results in loss of connections in the network structure. The remaining structure is the wireframe of the network where each wire forms the skeleton of individual vessels. Once created, we use each vessel’s skeleton to reconstruct its corresponding vessel, and measure its properties such as width, length, orientation, and distance from points of interest. Finally, we use each vessel as a mask and apply it to the tumor channel (mCherry) to locate and count the number of melanoma cells spreading along the abluminal surface of that vessel.

In order to capture the cells and the details of the vessel structures in the brain, a minimum 10X magnification is required. Consequently, the whole brain image, used as the input file, is tremendously large (180 MP and higher). Quantifying images with this size or higher, requires computational resources that are beyond what an average laptop offers. To make the tool accessible to everyone, we systematically break down each image into 100 equally sized tiles (where appropriate, the function adds proper symmetric padding to make the width and/or length divisible by 10), and perform vessel quantification within each tile in series. This not only makes the computations feasible on a laptop, but also results in a linear boost in computation time. Vessel quantification within each tile uses the local coordinates of the tile. To generate the results for the whole image, we aggregate the tile level measurements and translate them to the global coordinate of the original image. While this approach enables us to make the computations feasible on a wider range of devices, it should be noted that there is a caveat to this approach. While we are breaking down the images into smaller pieces (tiles), the vessels that span beyond a single tile, are cut and split between two adjacent tiles. This side-effect reflects in the results by introduction of additional (and shorter) vessels. In order to quantify the size of this side-effect, we counted the number of these vessels in different experiments, and noted that they account for 2-3% of the total vessels in the whole image. This range was small enough to safely disregard its effect on the final results. Vessel quantification is performed independently for each brain zone; i.e. tumor, control, and the remaining vessels. In the last step, the results from different zones are aggregated and the final results are exported as a CSV file, where each row represents a single vessel in the brain.

### Statistics

All analyses were performed using GraphPad Prism 5 (GraphPad Software). A p value of less than 0.05 was considered significant. All quantitative data represent the mean ± SD from at least 3 or more samples or experiments per data point. Additional experimental details (number of animals or cells and experimental replications) are provided in the figure legends. All box and whisker plots show the minimum and maximum values and a horizontal line at the mean.

## Author contributions

ACD and YZ designed and conceptualized the study and wrote the manuscript. YZ carried out the experiments. CPR assisted with *in vivo* and *in vitro* functional studies and wrote the manuscript. SW, SP and PK assisted with bulk RNAseq and RNA expression analysis. TB provided critical reagents or advice for carrying out portions of the study. SM provided critical feedback and edited the manuscript. AR and DK designed and helped implement the VeCo platform for high-throughput vessel analysis. All authors have been provided with a copy of the complete manuscript prior to submission.

## Competing interests

There are no competing interests to declare.

## Data availability

All sequencing data will be uploaded to NCBI GEO at the time of publication.

## Acknowledgements

ACD is supported by grants from the National Institutes of Health/National Cancer Institute (2RO1 CA177875 and RO1 CA2558451), the Melanoma Research Alliance (ID612638), and funds from the Emily Couric Cancer Center at the University of Virginia. SW is supported by an F30 from the National Cancer Institute (1F30CA268842-01) and a Melanoma Research Foundation Medical Student Research Grant. Portions of this research were supported by the NCI Cancer Center Support Grant 5P30CA044579 and by the UVA Genome Analysis and Technology Core (RRID:SCR_018883). Additional support was provided by The University of Virginia Flow Cytometry Core (RRID: SCR_017829). The Sony MA900 Cell Sorter was funded through the NIH S10 instrument program (S10 Grant Number 1S10OD028518-1). We would like to thank the UVA Biorepository and Tissue Research Facility (BRTF) for providing the human melanoma brain metastases sections. Special thanks to Constance Brinckerhoff and David Mullins for providing D4M melanoma cell line. And thanks to Brian Crompton for providing CRISPR/Cas9 construction vectors.

## Figures and legends

**Fig. S1.**
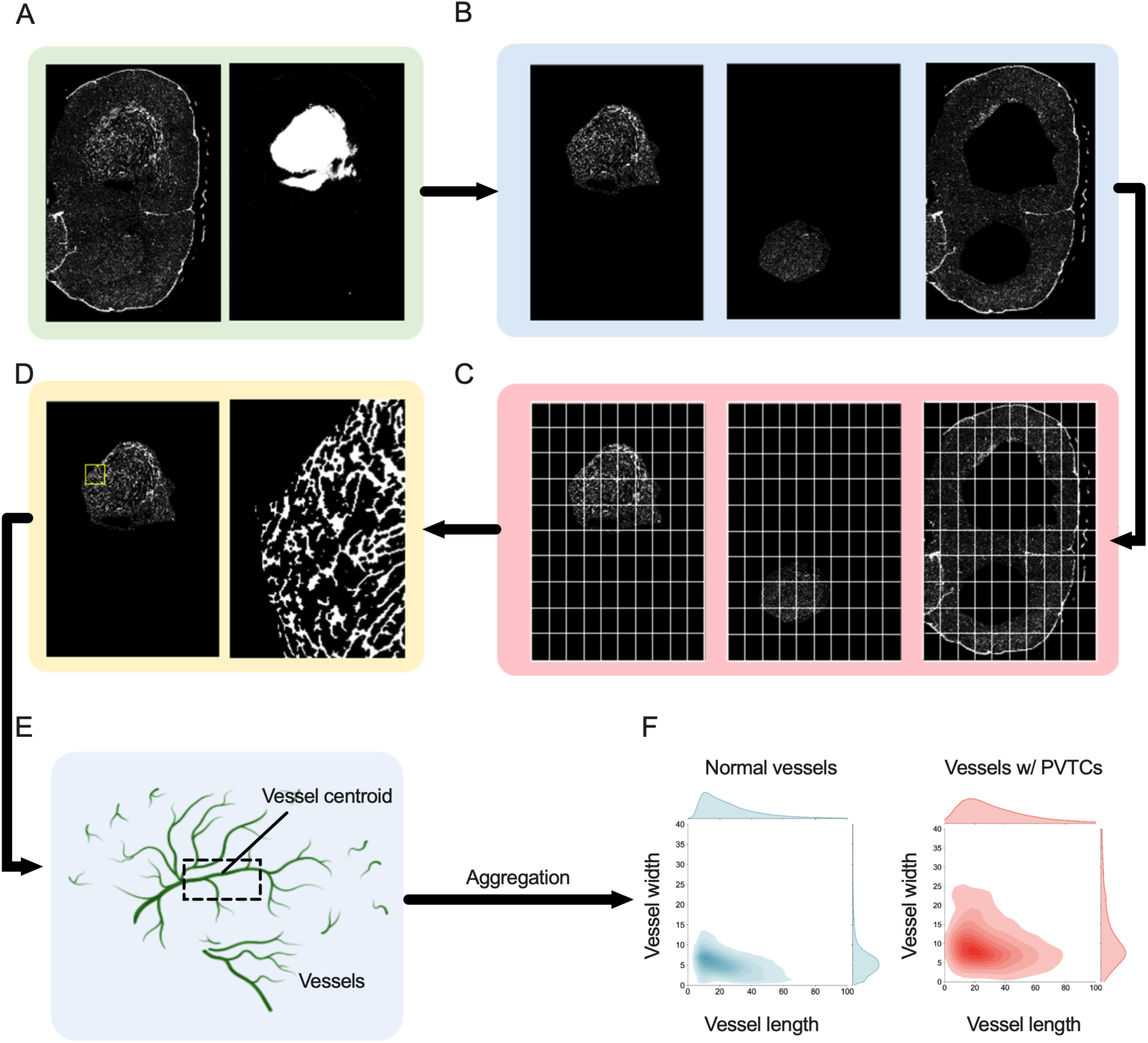
VeCo process chart. (A) Whole brain slice fluorescent images of blood vessels (ZSGreen^+^) or tumor cells (mCherry^+^) are imported to be analyzed (mCherry^+^ D4M tumors analyzed). (B) After pre-processing the images, three different zones (bulk tumor, contralateral striatum control, and remainder of the brain) are separated by cropping bulk tumor and control regions. (C) To make computations feasible on an average laptop, the images are broken down into 100 equally sized tiles to be processed in turn. (D) Object sizes are conserved in the breakdown process; however, each piece has its own coordinate system that should be translated to the global coordinate system of the original image after processing is finished. (E) Vessels are defined as the object between two joint points on the vessel’s network structure. Geometrical properties of vessels length, width, angle, and local position are quantified in each tile. (F) The quantified vessel measurements are aggregated within each tile, and then measurements from different zones of the brain are aggregated, compiled, and exported. Representive plots show length and width distribution of vessels with/without PVTCs.

**Fig. S2.**
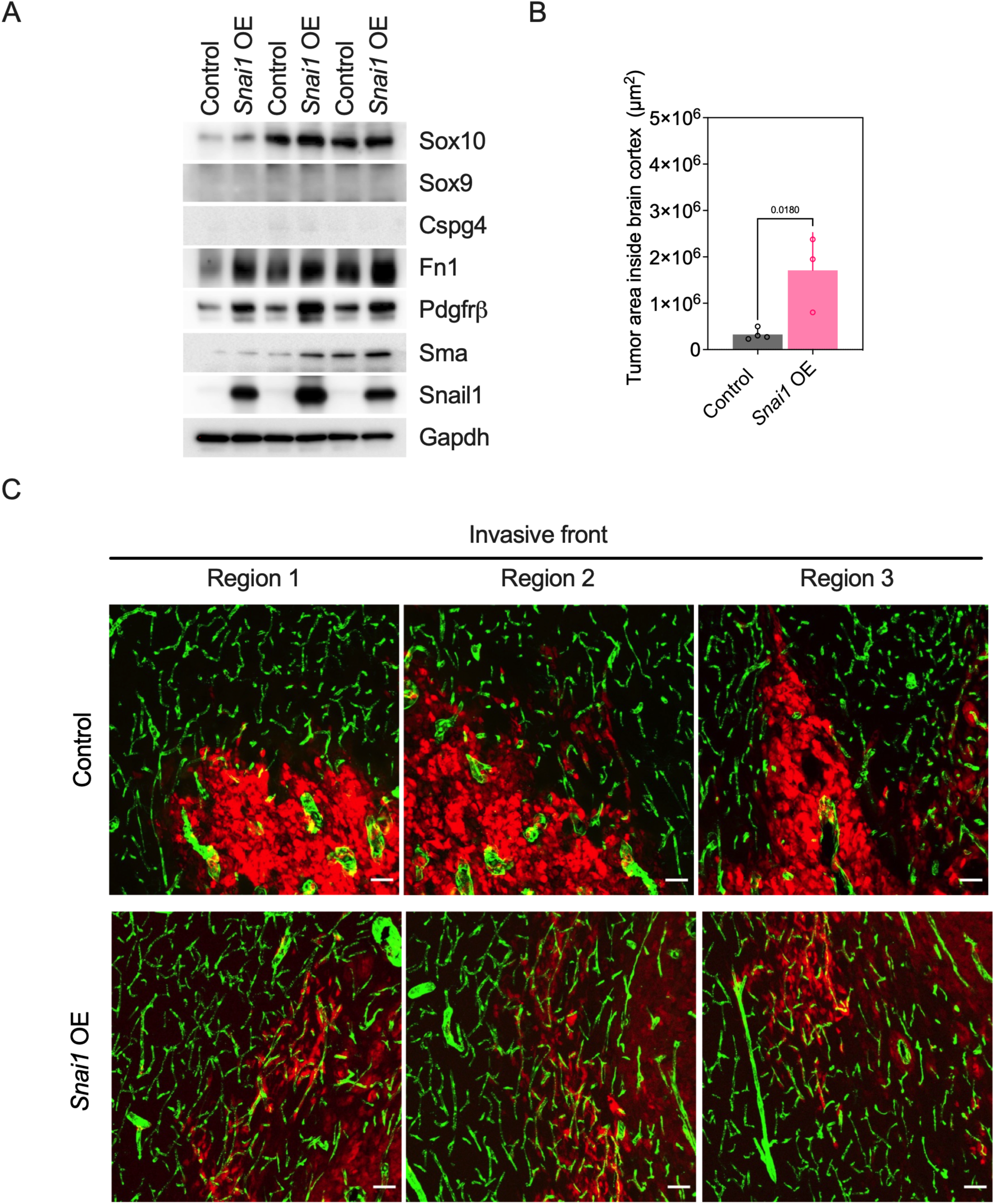
Overexpression of *Snai1* promotes perivascular invasion and tumor expansion in the brain. (A) Protein expression in B16F10 control or *Snai1* OE cells (three independent experiments are shown). (B) Quantification of tumor area inside the brain cortex from mice intracranially injected with B16F10 control or *Snai1* OE cells; control, n = 4 mice, *Snai1* OE, n = 3 mice. (C) Representative images of the invasive front in EC^iZSGreen^ mice injected with mCherry^+^ B16F10 control or *Snai1* OE cells. Scale bar, 50 μm (C). P values were determined by Student’s t-test.

## Notes

There are no conflicts of interest to disclose

### Competing Interest Statement

The authors have declared no competing interest.

### Summary of Updates

1) Author middle initials were added. 2) An additional Snail KO clone was added to figure 4.

